# Analyzing Nested Experimental Designs – A User-Friendly Resampling Method to Determine Experimental Significance

**DOI:** 10.1101/2021.06.29.450439

**Authors:** Rishikesh U. Kulkarni, Catherine L. Wang, Carolyn R. Bertozzi

**Affiliations:** Department of Chemistry, Stanford University, Stanford, California 94305, United States; Stanford ChEM-H, Stanford University, Stanford, California 94305, United States; Howard Hughes Medical Institute, Stanford University, Stanford, California 94305, United States

## Abstract

We report *Hierarch*, a Python package to perform hypothesis tests and compute confidence intervals on hierarchical experimental designs. Using a combination of permutation resampling and bootstrap aggregation, *Hierarch* can be used to perform hypothesis tests that maintain nominal Type I error rates and generate confidence intervals that maintain the nominal coverage probability without making distributional assumptions about the dataset of interest. *Hierarch* makes use of the Numba JIT compiler to reduce *p-*value computation times to under one second for typical datasets in biomedical research. *Hierarch* also enables researchers to construct user-defined resampling plans that take advantage of *Hierarch’s* Numba-accelerated functions. *Hierarch* is freely available as a Python package at https://github.com/rishi-kulkarni/hierarch.

## Introduction

Typical experimental design in the life sciences produces hierarchical data (or clustered, nested, multilevel, etc.)^1–3^ For example, a researcher might image multiple fields of view from the same coverslip in an imaging experiment or record multiple trials from the same animal in a behavioral study (**Scheme 1**). Despite the ubiquity of this type of experimental design, strategies for computing p-values for these experiments are hugely inconsistent in the literature. Common approaches range from “pseudoreplication” strategies that treat different fields of view as independent samples, to “summary statistic” approaches that aggregate the fields of view before performing a t-test or ANOVA.^4–6^ These approaches can produce wildly different p-values on the same datasets because they do not consider the hierarchical nature of the experimental design. The p-value is commonly misunderstood to be a measure of the compatibility of the null hypothesis with the observed data; however, the p-value is more accurately defined as a measure of the compatibility of the entire statistical model (including ALL assumptions made by the hypothesis test) with the observed data.^7^ If a researcher wishes to compute a useful p-value for a hierarchical dataset, the experimental design must factor into the statistical model in some manner.

**Scheme 1.**
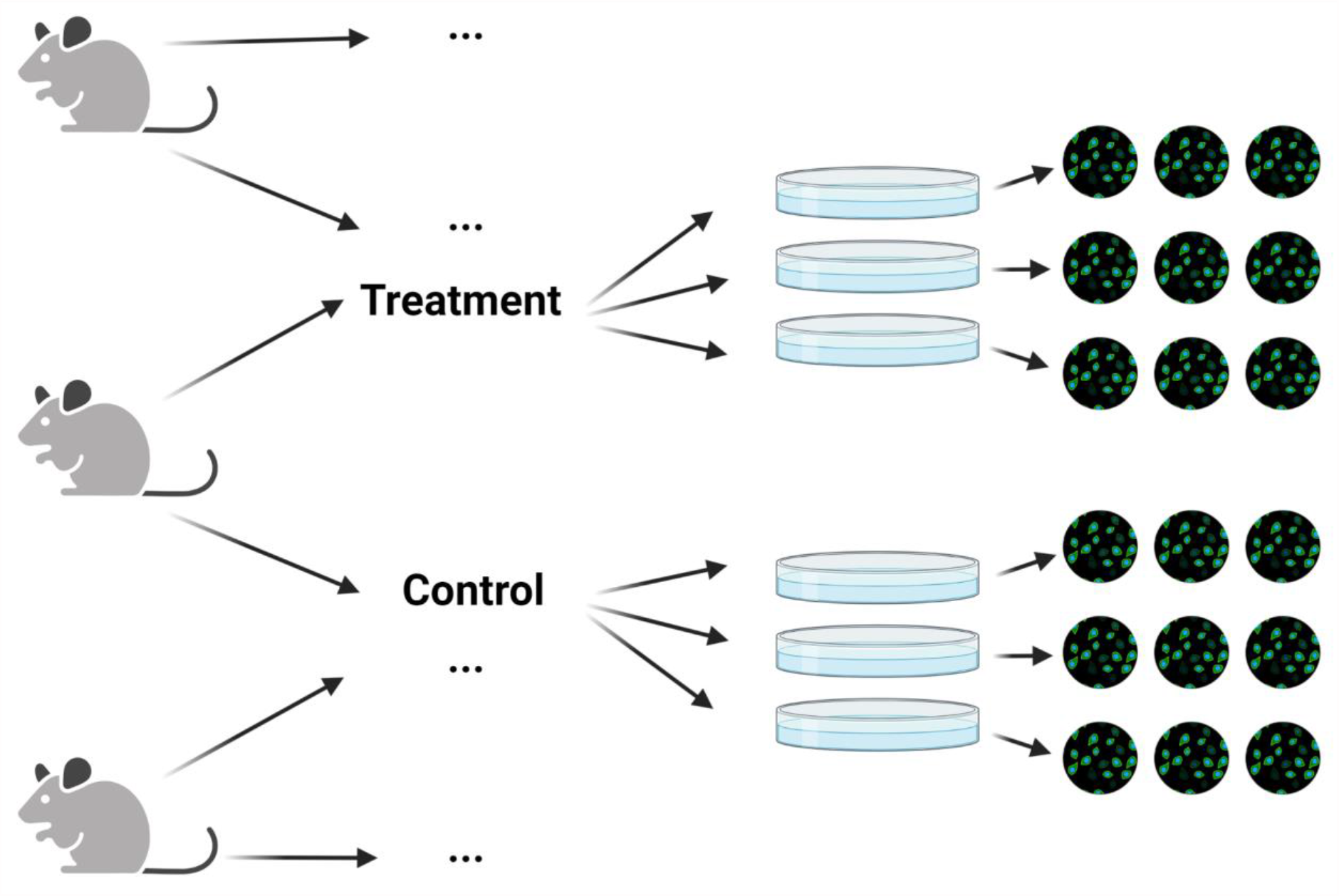
Hierarchical experiments are common in biomedical research. An example of a hierarchical experiment is an imaging experiment, where cells are isolated from donors, which are then treated in separate wells, which are then imaged under a microscope. These experimental designs are common in many fields of research, but especially so in molecular biology, imaging, and neuroscience.

One approach to analyzing hierarchical data is using a linear mixed model (or hierarchical model).^8,9^ Linear mixed models represent hierarchical data by being hierarchical themselves - the regression coefficients and intercept are themselves represented by another regression model. As flexible and powerful as they are, most studies employing linear mixed models involve very large numbers of clusters (>20), while studies in biomedical research typically have fewer than seven clusters and most often three to five.^10,11^ Simulation studies have shown that linear mixed models fail to control Type I error (false positive) rates with such a small number of clusters, becoming conservative or liberal depending if the effect of interest is within-clusters or between-clusters.^4,12^ Furthermore, the process of selecting parameters for a linear mixed model can be challenging – specifying the structure of a given data set is nontrivial, but failure to do so correctly completely invalidates the p-values computed by the model.

Ideally, researchers could analyze hierarchical data using a hypothesis test that incorporates data from every level of hierarchy, does not make any distributional assumptions about the dataset, and can be easily applied to a wide range of experimental designs. Randomization (or permutation) tests can be used to calculate p-values and confidence intervals while making only very weak assumptions about the nature of the data.^13,14^ By accounting for each level of hierarchy in the resampling plan, a hierarchical randomization test can control false positive rates while achieving good statistical power. Furthermore, resampling-based tests can be “distribution-free” in the sense that they typically make weaker assumptions regarding the population distributions underlying the samples.^15–17^ This has the added benefit of producing a p-value that does not depend on unverifiable assumptions about the data-generating process.^18^ Despite the good properties of resampling-based tests, they come with a few drawbacks. One major drawback of this approach is that for a given dataset, the script executing the resampling plan is often bespoke and computationally intensive. Furthermore, incorrectly specifying the resampling plan can result in inflated Type I error rates the same way that choosing the wrong traditional hypothesis test can inflate Type I error rates. Nonetheless, biological experiments often are by their very nature hierarchical, and demand a statistical approach that keeps hierarchy in mind.

To address these challenges, we present *Hierarch*, a Python-based module for hierarchical hypothesis testing. *Hierarch* is a lightweight Python module for nonparametric hierarchical bootstrapping and permutation testing based on NumPy^19^ and Numba.^20^ In this paper, we validate the Type I and Type II error rates of hierarchical randomization tests in *Hierarch* against asymptotic tests and walk readers through their usage. We compare the properties of these tests in simulation studies with the small sample sizes typical of biological experiments (n = 3 to 4 clusters), with different underlying population distributions, and with varying levels of hierarchy. We conclude that hierarchical resampling-based hypothesis tests are powerful, maintain better control of Type I error rates than asymptotic tests in a wide variety of conditions, and enable researchers to smoothly include multiple levels of clustering beyond the classic “biological replicate” and “technical replicate” dichotomy.

### How can you tell if your data is hierarchical?

Hierarchical data arises from one (or both) of two design issues (**Scheme 2**).^21^ The first issue to consider is hierarchical sampling, in which the sampled entities and the treated entities are not the same. For example, a researcher studying macrophages collects those macrophages by drawing a random sample of blood from a random sample of mice, then applying treatments to different wells in a 6-well place of the macrophages. The researcher has to account for the fact that random errors are introduced by *both* the mouse and the well – each mouse has different genetics and a different immune system, which introduces random errors to the measurement. Similarly, each well is delivered a slightly different number of cells and a slightly different amount of drug. Failure to account for both of these levels of hierarchy can result in unwarranted precision in the estimate of a treatment effect, which can fail to reproduce when the experiment is repeated in other mice.

**Scheme 2.**
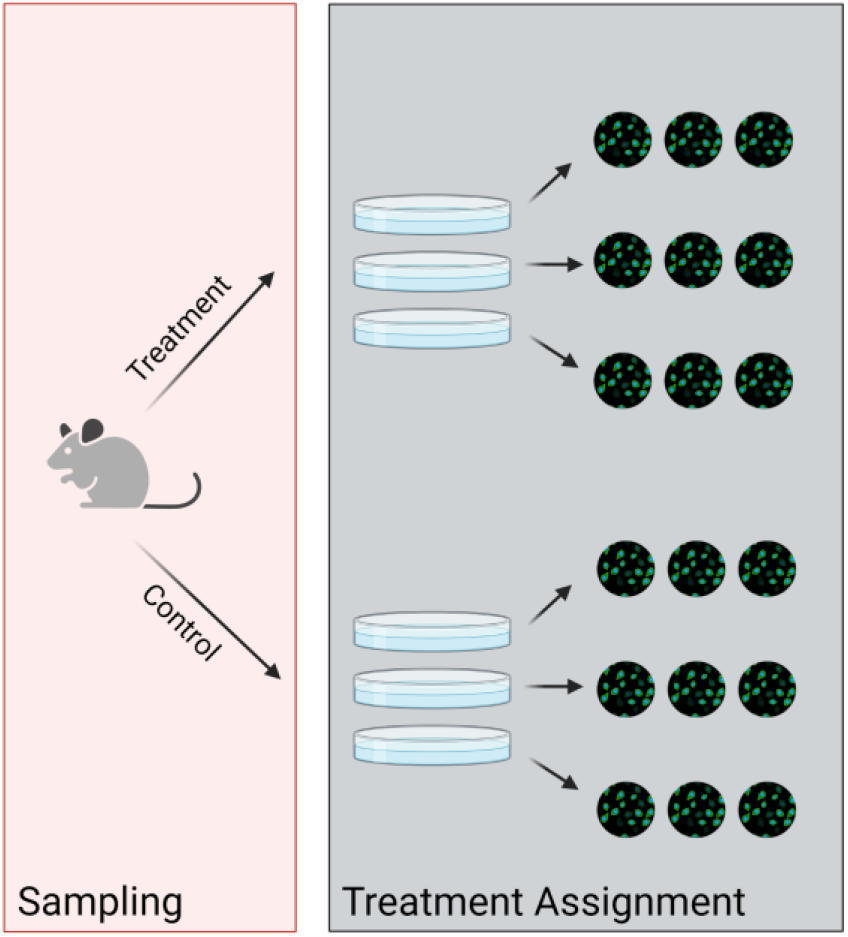
Hierarchy arises during sampling and treatment assignment. Hierarchy due to sampling occurs when the sampled entities and the treated entities are not the same. Hierarchy due to treatment occurs when the treatment entities and the observed entities are not the same.

The second design issue to consider is hierarchical assignment of treatment groups – or when the treated entities and the observed entities are not the same.^21^ For example, the researcher divides each mouse’s macrophages into six different wells and treats three of them with Treatment A, while the other three are treated with Treatment B. Then, the researcher performs an imaging experiment in which they look at several macrophages in each well. Because two macrophages in the same well are subjected to the same random errors in environment and treatment, they are much more similar than two macrophages in different wells. Again, failure to account for these levels of hierarchy can result an overly precise estimate of the treatment effect that disappears upon replication.

Under this framework, the vast majority of molecular biology and neuroscience experiments have at least three, if not four levels of hierarchy. Unfortunately, these design issues are difficult (and sometimes impossible) to avoid due to reasons of cost, ethics, or sample availability. However, by using statistical tools that understand hierarchical data, researchers can compute robust effect sizes that do not over- or under-estimate their confidence.

### Strategy for non-parametric analysis of hierarchical data

Permutation tests are a natural way to test the null hypothesis that a treatment has no effect, and the two samples are drawn from the same distribution, or the strong null hypothesis. Rather than using a theoretical null distribution, a permutation test builds a null distribution by shuffling the treatment labels in the data and recomputing the value of the test statistic. A permutation test assumes global exchangeability – that is, each observation was randomly assigned to one treatment or the other. Importantly, the null distribution in a permutation test is *only* conditioned on the observed data and the experimental design, so no unverifiable assumptions are made about the underlying data-generating process. For this reason, design-based permutation tests have been called the “platinum standard”^17^ of statistical analysis that ought to be given the “right to first refusal”^13^ when choosing an analysis for a given experiment. Permutation tests are computationally intensive; however, they have become more and more practical as personal computers have gotten faster.

Permutation tests face two key challenges when performed on hierarchical data. First, hierarchical data violates the basic assumption of global exchangeability.^22^ That is, while the labels of “treatment” vs. “control” are exchangeable under the null hypothesis, cells from different wells are not exchangeable. Again, this is because cells in the same well are subject to the same random errors at the well level and are expected to be more similar than cells from different wells. This problem can be avoided by only permuting on exchangeable levels (**Scheme 3a**). When analyzing experimental data, this means permuting the level at which the treatment was administered. This leads us to the second problem. When there are only a small number of available permutations and the researcher wishes to perform a two-tailed test, the empirical null distribution is too coarse for the p < 0.05 significance level. For example, with n = 3 in each group for a two-tailed hypothesis test, the smallest false positive rate that can be achieved is 0.1. At n = 4 per treatment, the only alpha below 0.05 is 0.028. Only at n = 5 per treatment or more can the experimenter control alpha at values close to 0.05. We note that the most robust way around this issue is to perform experiments with at least n = 5 per treatment. However, it is sometimes impossible to acquire more samples, for example in cases where samples are sourced from human subjects. Ordinarily, this leaves the researcher stuck between a rock and a hard place - either they have to go with the strong assumptions of an asymptotic test (which, at n = 3 per treatment condition, are doing at least as much work as the data is) or accept that they cannot achieve p < 0.05 with a nonparametric test.

**Scheme 3.**
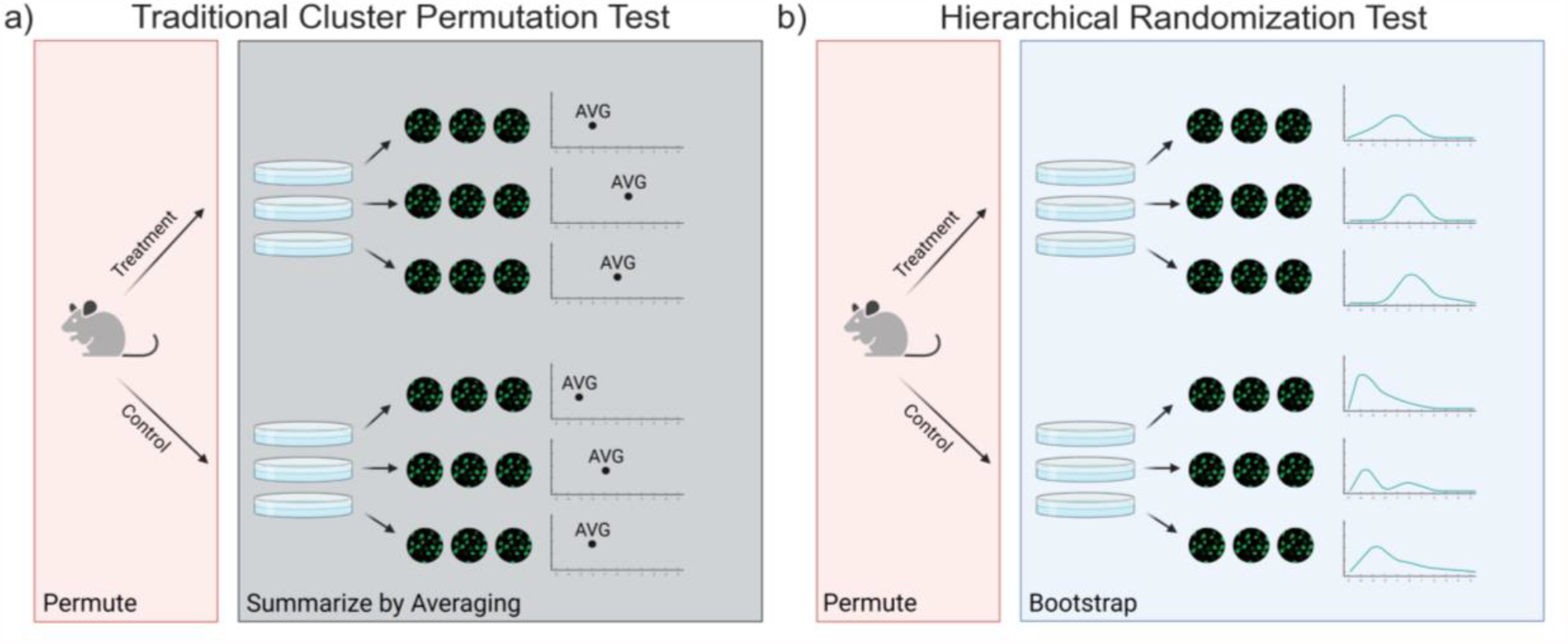
Hierarchical randomization combines permutation and bootstrapping to perform a hypothesis test. a) By averaging, traditional cluster-based permutation tests shuffle discard information from the levels of hierarchy that arise due to treatment assignment. b) Hierarchical randomization tests use bootstrapping to compute posterior distributions of the mean for each treated sample, thereby using all of the data collected to compute a p-value.

In this example, a traditional cluster permutation test would involve summarizing the observations in each well by taking the average, then permuting the treatment labels to form a null distribution. With only 6 total wells, however, there are only 20 possible permutations, so the minimum two-tailed *p*-value that can be computed is 2/20, or 0.1. Instead, we propose a test that shuffles posterior distributions of the cluster means rather than merely the point estimates of the cluster means (**Scheme 3b**). To estimate these posterior distributions, we utilize another resampling-based method. The nonparametric bootstrap, developed by Efron^23^ and extended by many others,^2,24–26^ is an attractive method to nonparametrically estimate the posterior distribution of each cluster mean in this situation. The bootstrap procedure involves resampling the within-cluster observations with replacement and recomputing the mean many times (say, 1000), resulting in a distribution of means that, importantly, reflect the standard error and skew of the original observations. Then, each set of bootstrap means is shuffled some number of times (in this case, 20) and the test statistic is recomputed with every shuffle. The *p*-value is the fraction of these t statistics that are as or more extreme than the observed t statistic. This strategy, which is performs bootstrap aggregation of several permutation tests,^27^ enables researchers to incorporate the observed within-cluster variability into the hypothesis test and control alpha at 0.05 for datasets with as few as 6 clusters.

### Description of the hierarchical randomization test

To explain our algorithm, we consider again the above dataset consisting of two treatments, three wells each, and three images each (**Figure 1a**). The researcher seeks to test the null hypothesis that the treatment had no effect on the mean fluorescence intensity of each image.

**Figure 1.**
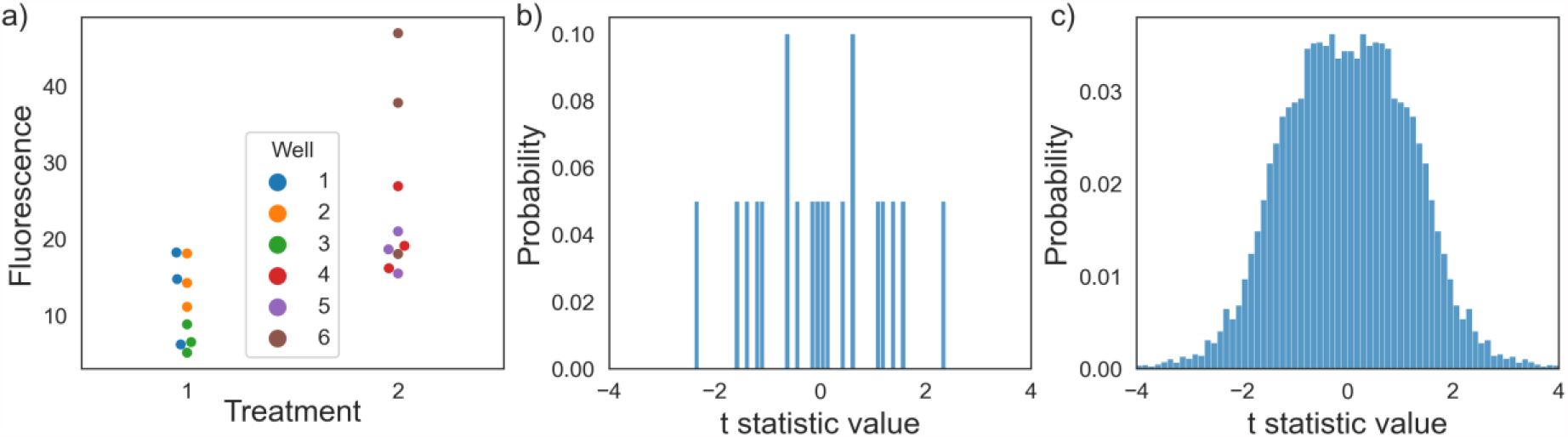
Hierarchical randomization uses bootstrapping to construct the empirical null distribution. a) Simulated data corresponding to an experiment in **Scheme 3**. b) A traditional cluster permutation test would only have twenty possible permutations, resulting in the smallest calculable two-tailed p-value being 0.1. c) Hierarchical resampling constructs a full empirical null distribution, resulting in a p-value of 0.0429.

1. First, calculate the observed value of the test statistic. For a difference-of-means test, we use Welch’s t statistic.
2. Then, for each biological replicate, draw a bootstrapped sample (resampling with replacement) from its technical replicates.
3. Then, permute the “treatment” labels and calculate a test statistic. Repeat this step a number of times (in this example, 20 times).
4. Repeat steps 2 and 3 a large number of times (>500) to generate the empirical test statistic distribution.
5. Determine what fraction of the empirical test statistic distribution is as or more extreme than the observed test statistic. This number is the two-tailed *p*-value.

A traditional cluster permutation test would only be able to produce a null distribution containing at most 20 possible values (**Figure 1b**), but hierarchical randomization generates a full null distribution without making distributional assumptions about the data (**Figure 1c**). More generally, the algorithm deals with each level of hierarchy in one of two ways. For hierarchy arising due to treatment assignment, the algorithm uses nonparametric bootstrapping to estimate the sampling distribution of the mean. For hierarchy due to sampling, the algorithm restricts the number of possible permutations such that only “within-cluster” permutations are possible. This procedure mimics the data-generating process under the null hypothesis (that is, the hypothesis that the treatment did nothing at all). First, each well is resampled from its fields of view and then randomly assigned to one of the two “treatment” labels. Using *Hierarch*, this procedure is fully automatic - once the researcher has specified their experimental design by organizing their data, the algorithm will produce a *p*-value without requiring any further input. Moreover, the algorithm infers the correct resampling plan for any hierarchical experimental design. If a researcher pre-commits to using hierarchical randomization as their analysis tool of choice, they have eliminated an important researcher degree of freedom - choice of hypothesis test - whilst retaining the flexibility to analyze a wide range of experimental designs.

### Caveats

The most important assumption this test makes is that the labels being shuffled are exchangeable under the null hypothesis. In other words, it assumes that the clusters attached to the labels were assigned randomly. The second, weaker assumption is that the observations have similar distributions (though not necessarily normal). This is a weak assumption because by using an approximately pivotal test statistic (such as the t statistic),^16,28^ the assumption of homogeneity of variances does not have to be fulfilled for this test to maintain control of Type I error rate. However, with very few clusters, this test can be sensitive to heterogeneity of variances (see simulation study below).

An important consideration with this approach is that bootstrapping is only appropriate when the within-cluster data points represent a random sampling of possible within-cluster values. For example, an imaging experiment might involve taking images of several fields of view within a well and measuring some per-cell quantity in each image. In this case, the fields of view are randomly sampled from all fields of view in the well (as there are fields of view that were ignored) and therefore can be resampled, but cells within each field of view are not randomly sampled (as every cell in a given field of view is measured) and therefore should not be resampled via bootstrapping. For a deeper discussion of this, see van der Leeden, et al.^9^

Another consideration to using these resampling techniques is that the permutation test is no longer exact - there are usually a much larger number of possible resamples than can reasonably be calculated (though in this simple case, it is possible). However, by performing a large number of resamples and using an appropriate test statistic, this approximate test will have size close to 0.05 and good power while not requiring the researcher to make distributional assumptions about their dataset. To demonstrate the flexibility of hierarchical randomization tests, we will discuss the analysis of three datasets.

### Example 1: Three-Level Mouse Socialization Experiment

First, we consider the behavioral assay shown in **Figure 2a**. Here, the researcher has a control group and treatment group of four mice each. Each mouse performs 500 trials of a behavioral assay to test the hypothesis that the treatment causes an increase in socialization duration. The multilevel design (treatment -> mouse -> trial) of this experiment lends itself to a hierarchical hypothesis test. Furthermore, given that the units of the measurement (seconds) are bounded by zero, we have good motivation to try a test that does not assume normality. We consider the following data-generating model:

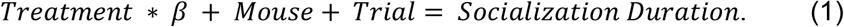

**Figure 2.**
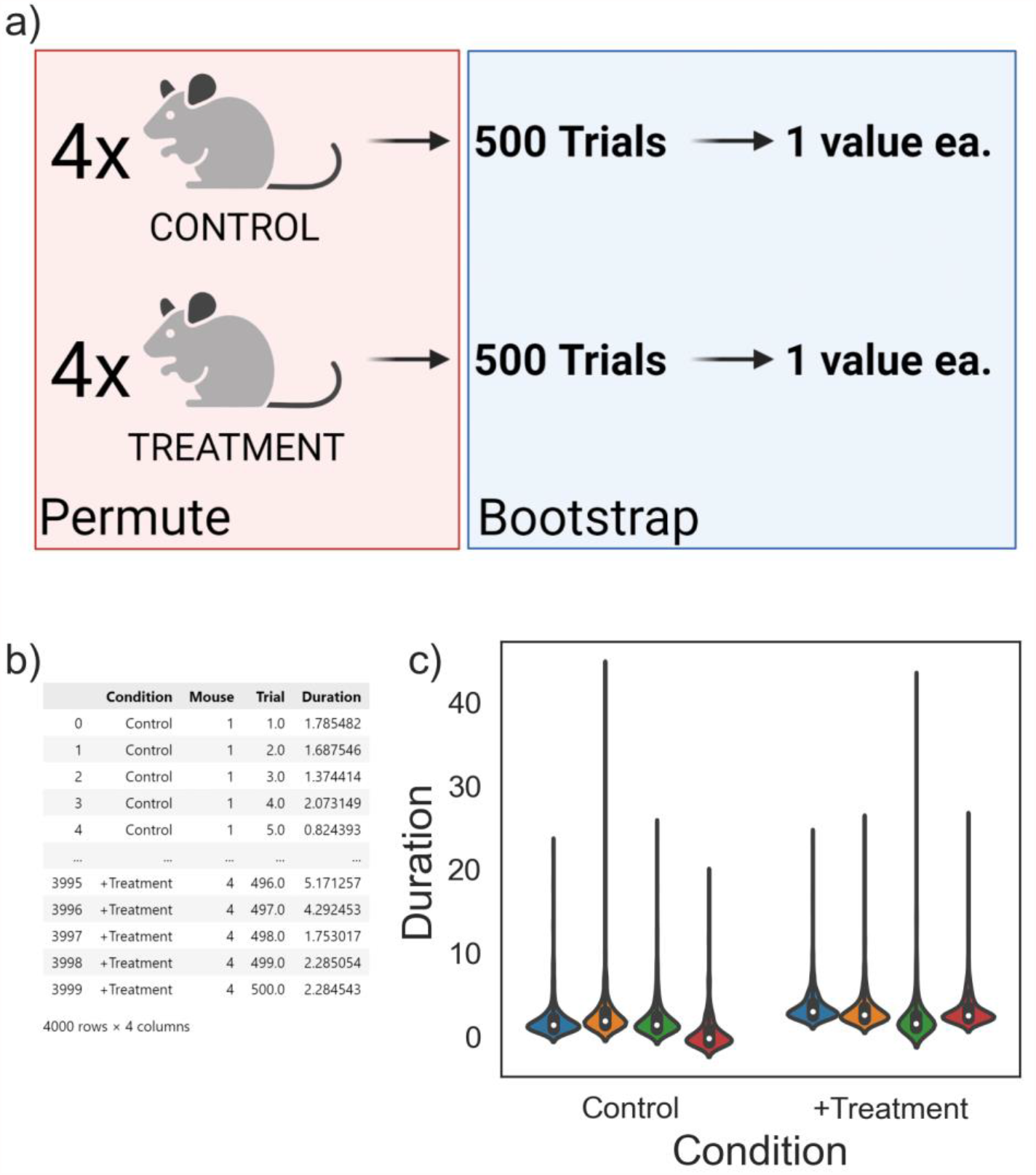
Using hierarchical randomization to analyze a mouse behavioral study. a) Simulated data corresponding to an experiment with two treatments, four mice each, 500 trials each. b) A table of raw data collected in this study. By organizing the input data into columns corresponding to the experimental design, *Hierarch* can automatically infer the correct resampling plan for the dataset. c) Violin plots illustrating the skewed nature of the dataset. p = 0.037, hierarchical randomization, p = 0.038, Student’s t test.

The researcher seeks to estimate the treatment effect *β* and calculate a p-value against the null hypothesis that the treatment effect is 0. Each mouse, however, has an individual random constant that reflects mouse-to-mouse variation in baseline socialization duration. While the terms in the model can be written in any order, it can be helpful to structure the equation in the same order as the actual hierarchical experiment. In this case, treatments are assigned to mice, which are measured 500 times each. Simulated data for this experiment is shown in **Figure 2b, c** with a true effect size of 1 second. The researcher can organize their data into the “long” format presented in **Figure 2b** (visualized in **Figure 2c**), where each column corresponds to one of the terms in the model (treatment, mouse, trial number). Given that the input data is organized such that the column order mimics the experimental design, *Hierarch’s* hypothesis_test function will conduct the appropriate resampling plan by default.

According to the hierarchical randomization algorithm, treatment labels are permuted only at the “mouse” level - individual behavioral trials are never exchanged between different mice. This ensures that the test does not break the dependence structure that exists in the dataset. Instead, uncertainty in the mouse-level mean for the behavioral trials is represented via bootstrapping. *Hierarch* performs 35,000 resamples (500 bootstraps with 70 permutations each) in less than 200 milliseconds and generates a two-tailed *p*-value of 0.037, indicating a statistically significant difference. In this case, we note that the hierarchical randomization *p*-value is similar to the *p*-value calculated by a two-sample t-test after averaging the 500 trials for each mouse (*p* = 0.038). Given the large number of trials, the standard error of the mean for each mouse is quite low, so computing an average socialization duration for each mouse does not throw away much information. Rather, we use this as an illustrative example to show that even when another test might be appropriate, hierarchical randomization produces a similar *p-*value, but can be applied to more complicated datasets without much trouble.

While *p*-values are useful, they are best paired with a measure of effect size.^29,30^ Generally speaking, accompanying the effect size with a confidence interval gives readers more information with which to interpret experimental results. One trouble with computing confidence intervals for effect sizes is that few methods actually maintain the nominal coverage when sample sizes are small – 95% confidence intervals often do not contain the true value exactly 95% of the time.^31–33^ Hierarchical randomization tests can be inverted to form confidence intervals that are very close to exact. This is a key advantage over the t-test for non-normal data – t-intervals, which are quite robust for small samples, tend to be conservative for non-normal data and will produce too-wide confidence intervals. As we will show in the simulation study below, the 95% confidence intervals produced by *Hierarch’s* confidence_interval function do indeed contain the true effect size 95% of the time, even for datasets with as few samples as this one. In this case, *Hierarch’s* 95% confidence interval on the effect size is (0.179, 2.697) while the corresponding t-interval is (0.0166, 2.607). This is an interval around the beta coefficient in equation 1, which we simulated with a value of 1.

### Example 2: Four-Level Imaging Experiment

Next, we consider the motivating example from above: a paired experimental design common in molecular biology and neuroscience. Here, the researcher is interested in testing the effects of a drug on neuronal firing rate in a culture model. From each of three pregnant mice, the researcher prepares six separate neuronal cultures in a six-well plate. In each plate, the researcher treats three wells with the drug of interest and gives three a vehicle control. Then, the researcher performs a current-clamp experiment to measure the firing rate of three neurons in each well (**Figure 3a, b**). We consider a data-generating model as follows:

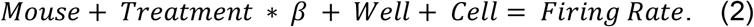

**Figure 3.**
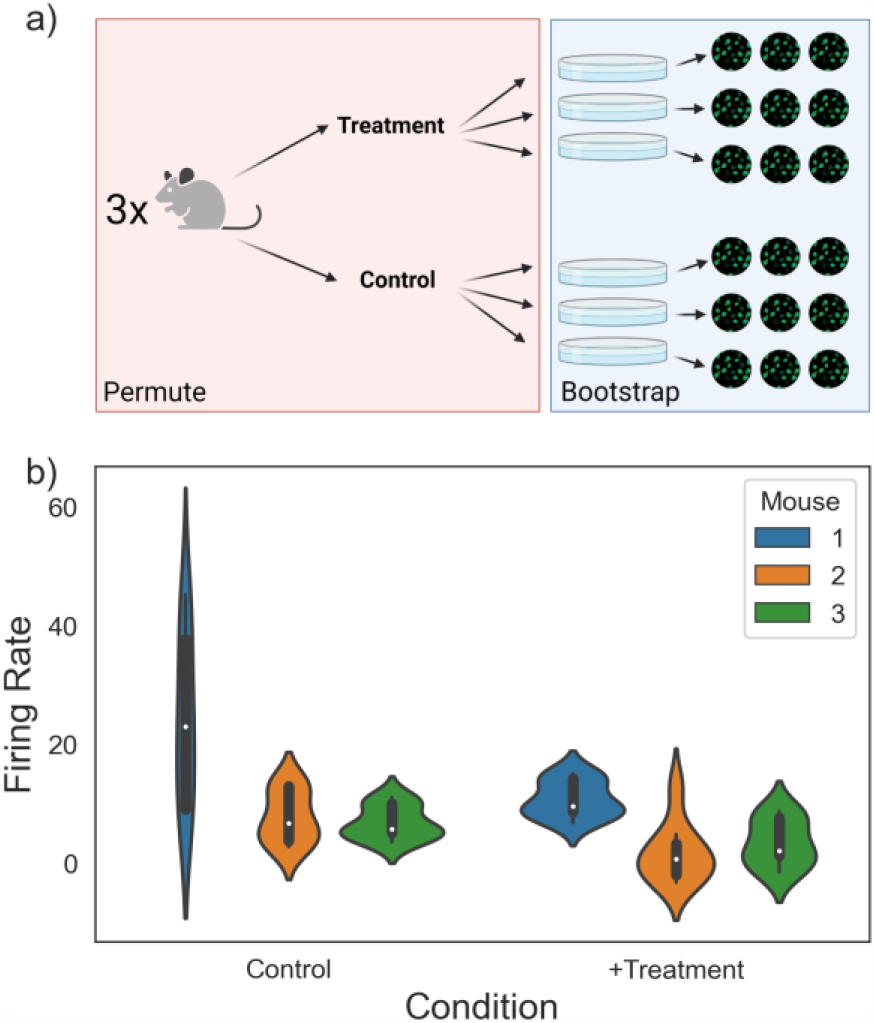
Analyzing a four-level imaging experiment. a) An experiment with three mice, two treatments each, three wells each, and three neurons each. b) Violin plots visualizing the dataset. p = 0.027, hierarchical randomization, p = 0.101, pooled Student’s t test.

The researcher seeks to estimate the fixed treatment effect *β* and calculate a p-value against the null hypothesis that *β* is 0. We simulated the data in **Figure 3b** with an effect size of 11. Unlike the previous experiment, there is an additional constant term - we assume that not only does each well have a random baseline, but each mouse *also* has a random baseline. Despite how common this experimental design is, it is not immediately clear how best to calculate a p-value with a traditional approach. Should the researcher perform a Student’s t-test with n = 9 wells in each treatment group? If each mouse has a different baseline firing rate, however, the between-mouse variance would erode the power of the test. Furthermore, the t test assumes that the treatment effect is fixed and neglects the fact that, at least on one level, the data is paired. On the other hand, aggregating the firing rates up to the treatment level and performing a paired t test with n = 3 also has little power by virtue of reducing the sample size to 3.

Two traditional options can be used in this situation: either treating each mouse as a separate experiment and combining the data in a manner analogous to an individual participant data meta-analysis or fitting a mixed effect model.^34–36^ Both of these approaches require researchers to make distributional assumptions about their datasets, however. Hierarchical randomization provides a natural way to test a single hypothesis and generate a single p-value on the combined experiments - bootstrap the mean firing rate for each well from its neurons, then permute the treatment labels on the wells *within* mice.^37^ In this example, there are many more possible permutations (20^3 = 8,000), so the researcher can choose to run a subset of them (100 bootstraps, 4,000 permutations each). This results in a *p-*value of 0.027 and a 95% effect size confidence interval of (1.157, 14.916), which contains the true value of 11. Pooling all of the data and performing a t test gives a *p*-value of 0.101 and a 95% confidence interval of (−2.009, 17.305). By accounting for sampling hierarchy, hierarchical randomization can be more powerful than other non-meta-analytic approaches.

Given that we have assumed a fixed treatment effect, the experiments (mice) are automatically weighted by their sample size. However, hierarchical randomization makes it simple to analyze data from a random treatment effect perspective, as well. If the researcher suspects there may be significant heterogeneity in treatment effects - perhaps the donor mice are heavily outbred, or the donors are humans – they can incorporate this heterogeneity as an interaction effect (**Figure 4a**).^38^ The equation can be written as follows:

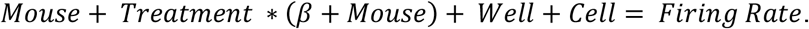

**Figure 4.**
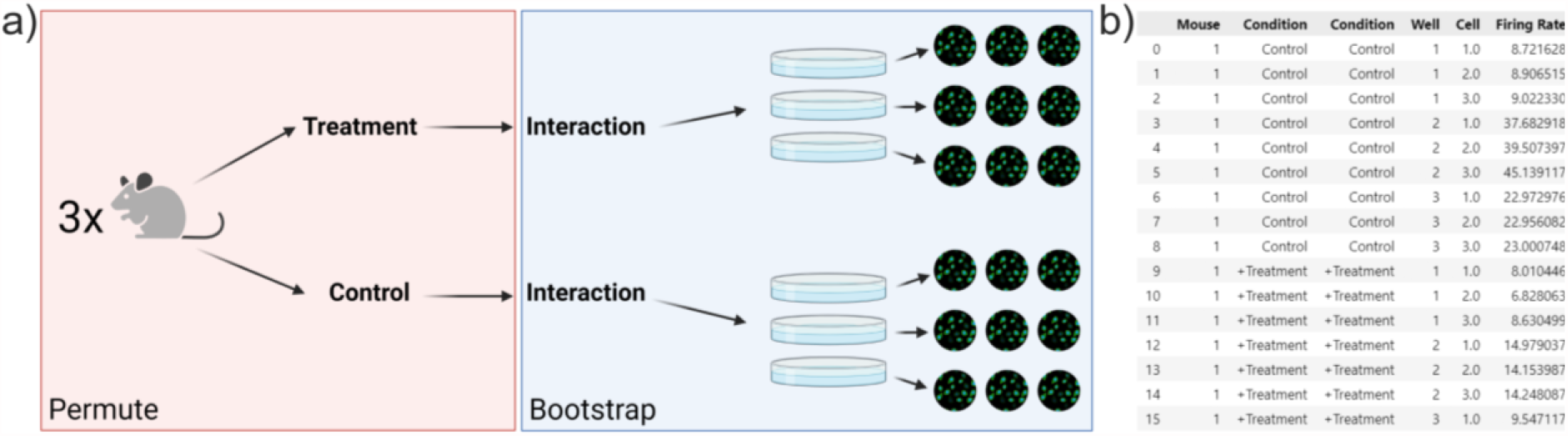
Incorporating heterogeneous treatment effects. a) Adding a mouse-treatment interaction term to the experiment design describes heterogeneity of treatment effects. b) By duplicating the “Condition” column in the input data, *Hierarch* will account for this interaction.

The *Mouse* term in the interaction is not the same value as the *Mouse* baseline constant – rather, it represents the fact that in this model, we are allowing each mouse to have a unique slope for the treatment effect. Distributing this equation gives:

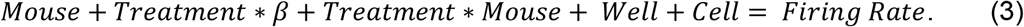

This equation splits the treatment effect into two terms – an average treatment effect (Treatment x β) and a random interaction effect (Treatment x mouse). Updating the raw data to include this additional term is all that is necessary for *Hierarch* to carry out the appropriate resampling plan for the random treatment effect model. This is most easily done by simply duplicating the “treatment” column in the raw data (**Figure 4b**), which communicates to *Hierarch* that an interaction term is present.

Accounting for treatment effect heterogeneity increases the *p-*value (0.118) and widens the confidence interval (0.478, 23.45, 90% confidence interval). This makes sense in the context of a random-effect model - in order to make a precise estimate of effect size, both the “within-mouse” sample size and the number of mice must be large. Performing a large number of samples within a single mouse may yield a very precise estimate of the effect size in that mouse, but if the effect varies mouse to mouse, the overall average effect can only be accurately estimated by studying several mice. Furthermore, when assuming random treatment effects, reporting a confidence interval on the effect size is more important than ever because the average treatment effect is entirely dependent on the mix of donors. Given that, summarizing the effect size with a single point estimate is too reductive – after all, there is no single number that can describe the true effect of the drug.

### Example 3: Four-Level Rat Behavioral Study with Multiple Time Points

Finally, we consider an experimental design with several treatments that seeks to test a single hypothesis. A researcher is interested in measuring changes in a neural population over the course of learning a task. The researcher has four rats, who are each measured on four days in the learning process. On each day, the rat attempts the task 500 times, during which some population of neurons is recorded via electrodes implanted in the rat’s skull. From each recording, the researcher computes some neural population-level metric (**Figure 5a, b**). The researcher considers the following model:

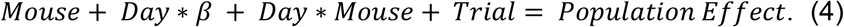

**Figure 5.**
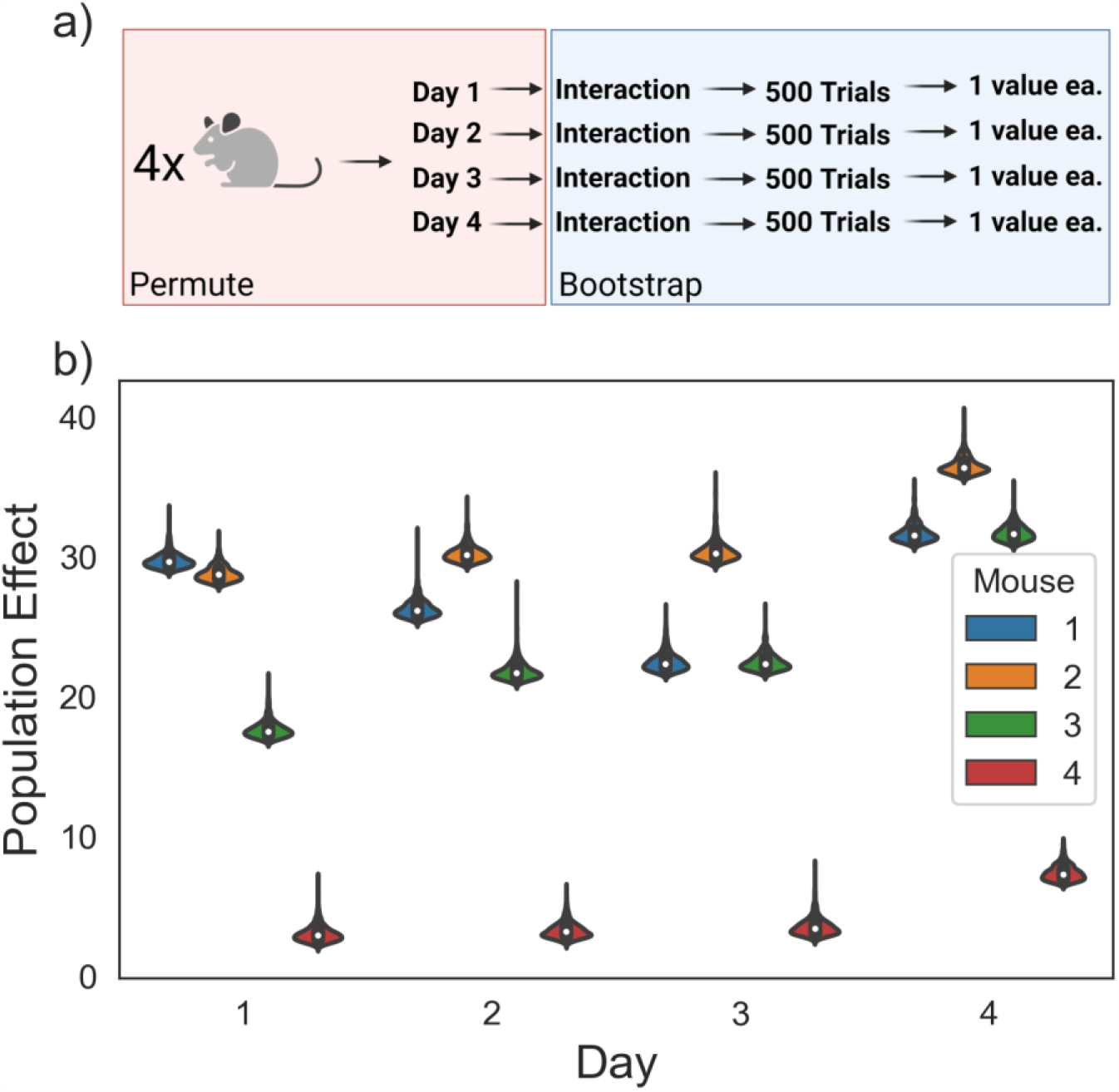
Analyzing an experiment with multiple treatment conditions testing a single hypothesis. a) An experiment with four mice, two treatments each, three wells each, and three neurons each. b) Violin plots visualizing the dataset. *p* = 0.024 (hierarchical randomization) for the hypothesis that there is a day-to-day increase in the population effect.

We include an interaction effect to account for day-to-day variation in the population effect due to electrode drift and other changes in the mouse that are unrelated to the task at hand. As above, we want to perform a hypothesis test against the null hypothesis that β = 0.

The experimental design poses another challenge, however - there are four different days. One approach could be to perform several two-sample tests between each day and the next day. However, this approach only considers a subset of the dataset at a time, and as a result loses a lot of power – none of the day-to-day comparisons are significant. Upon closer examination, this logic behind this approach is unclear – if we have a single hypothesis (that β ≠ 0), why perform multiple hypothesis tests? Another option is fitting a simple linear regression or a mixed model – but we have no reason to think the errors of the neural population measure are normally distributed and, as usual, we do not have many clusters. This example motivates the construction of a new studentized covariance test statistic that can be used to perform a single hypothesis test against the null hypothesis that β = 0 when there are multiple treatment groups with a hypothesized linear relationship.

This test statistic can be calculated on every shuffled dataset in a hierarchical randomization test, which provides a test against the null hypothesis that the slope for a given regressor in a linear model is equal to zero. For two-sample datasets, this test statistic has a linear relationship with the t-statistic and therefore will calculate the same *p*-value, which is demonstrated in the simulations below. In this instance, hierarchical randomization computes a *p*-value of 0.0236 and a 95% confidence interval of (0.42, 3.622), which contains the true, simulated value of β = 2.

### Construction of a studentized covariance test statistic

When constructing a randomization test for some parameter, the test is only guaranteed to be exact for the null hypothesis of the distributions being equal. To maintain a Type I error rate of 5% for the more general null hypothesis that the parameter is equal, the test statistic must be at least approximately pivotal – that is, its distribution does not depend on unknown parameters, such as the population standard deviation. Approximately pivotal statistics can be constructed following the procedure of Janssen,^16^ which was expanded by Chung and Romano and others.^28,39–41^ This is done by dividing the comparison of interest by an estimate of its standard error. As an illustrative example, we will discuss the construction of the t statistic. We consider a linear equation describing a data-generating process.

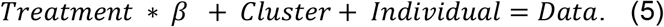

When there are only two treatment groups, β can be estimated using equation 6,

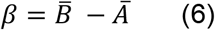

▪ 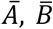 are the means of group A and B, respectively.

Student’s t test is a hypothesis test against the null hypothesis that β = 0. The t test is based on the t statistic, which is given by the following equation,

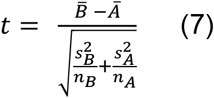

▪ 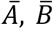 are the means of group A and B, respectively.
▪ 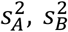 are the sample variances of group A and B, respectively.
▪ *n*_*A*_, *n*_*B*_ are the number of samples in group A and group B.

This is equivalent to the following expression:

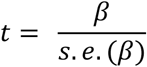

▪ *β* is the estimator for the slope in equation 5.
▪ *s. e*. (*β*) is the standard error of the estimator *β*.

This is a general approach for constructing an asymptotically normally distributed test statistic (or a Wald-like statistic). Because of this property, when a Wald-like statistic is used as the test statistic in a randomization test, the test gains asymptotic validity against unequal variances between treatment conditions and gives the researcher the ability to make directional conclusions. However, the t statistic can only be used as a test of β = 0 when there are only two samples. Instead, we can express β as a ratio between the covariance of X and Y and the variance of X:

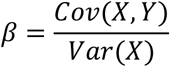

▪ *X, Y* are the treatment condition and observed data, respectively.

In a randomization test, we are merely shuffling the relationship between X and Y. Therefore, the variance of X is constant during the shuffling procedure. We can therefore construct a Wald-like test statistic for β using the covariance of X and Y, which is based on the work of DiCiccio and Romano:^42^

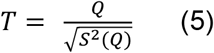

▪ *Q* is the sample covariance of *X* and *Y*.
▪ 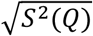 is standard error of the sample covariance of X and Y, or the square root of the sample variance of the sample covariance of X and Y.

The sample covariance of *X* and *Y, Q*, is given by equation 6:

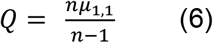

▪ *n* is the number of total observations.
▪ *µ*_*1,1*_ is the population covariance of *X* and *Y*, otherwise known as the first product central moment of *X* and *Y*. This is computed with equation 7:

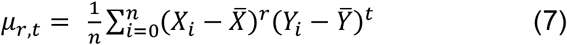

To compute the sample variance of *Q*, it is helpful to start with the population variance. For any distribution with defined moments, the population variance of the sample covariance is expressed by equation 8:

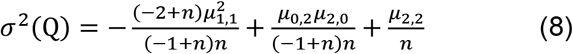

▪ *n* is the number of total observations.
▪ *µ*_*r,t*_ represent product central moments of *X* and *Y* given by the equation 7.

Equation 8 represents a biased estimator for the variance of *Q*, however. The unbiased estimator for the variance of Q, or the sample variance of Q, is prone to numerical instability (**SI Appendix 1**), so instead, we use the following bias-corrected approximation for the sample variance of Q,

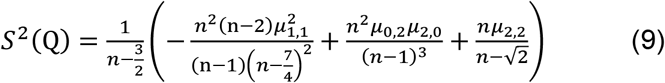

▪ *n* is the number of total observations.
▪ *µ*_*r,t*_ represent product central moments of *X* and *Y* given by the equation 7.

Using equation 9 as an estimator for the variance of the sample covariance of X and Y, we can use the Wald-like statistic in equation 5 as the basis of a hierarchical randomization test. In the simulation study below, we will investigate the properties of this test against nonnormality and heteroscedasticity.

### Simulation Study

In this section, we demonstrate that hierarchical randomization successfully controls Type I error rates without being sensitive to the underlying distribution of the dataset. We were particularly interested in small studies typical in biomedical research, so we chose to consider experiments structured similarly to the case studies detailed above. For larger datasets, there are numerous other simulation studies in the literature demonstrating the good properties of randomization tests.^13,28,41^

## Methods

All hierarchical randomization tests were performed using *Hierarch* version 1.1.1 (https://github.com/rishi-kulkarni/hierarch). All simulation code is available at https://github.com/rishi-kulkarni/hierarch-simulations. For Type I error rate control studies, p-values for t tests and linear regression were computed using scipy.stats. For confidence interval simulations, parametric confidence intervals were generated using the statsmodels package. We performed between 20,000 and 100,000 resamples per test depending on the total number of available permutations. The largest tests (100,000 resamples) took less than 200 milliseconds each. In each test, we set alpha to 0.05 and simulated 10,206 datasets (729 on each of 14 cores of an Intel i9-9940X CPU) according to one of the two following data-generating models:

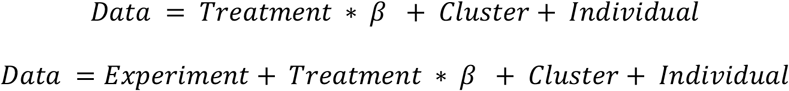

In each simulation, we generated the cluster baseline and the individual values with either normal, lognormal, Pareto, or gamma random variables. To demonstrate the general applicability of hierarchical hypothesis testing, we varied the ratio of within-cluster variance to total variance (intraclass correlation). The results of these simulations were evaluated for Type I error rate control – essentially, what percentage of the simulated datasets and hypothesis tests returned a *p*-value below 0.05 when there was no true difference between the datasets?

Confidence intervals are calculated using the test inversion procedure discussed in Manly.^43^ Briefly, the bounds of the hypothesis test’s rejection region are found using an iterative approach. These bounds are then unstudentized back to the units of the β coefficient. Each iteration was allowed to perform between 1,000 and 10,000 shuffles with a maximum of 10 total iterations per bound. These simulations were repeated for varying values of β. The results of these simulations were evaluated for coverage probability – essentially, what percentage of the 95% confidence intervals contain the true value of β?

## Results

First, we examined both types of hierarchical randomization tests (using the t statistic and using the studentized covariance statistic described in equation 5). We simulated datasets with two treatment groups in which the effect size (*β*) was set to zero (**Figure 6**). For each set of simulations, the fraction of hypothesis tests that returned a significant result was plotted on the *y*-axis. The shaded region represents the 95% confidence interval around a 5% Type I error rate, which each test should ideally remain in. As expected, the Student’s t test is only able to robustly maintain a 5% Type I error rate when the underlying data is normally distributed, while Welch’s t test is conservative in all cases. We observed that when there are 8 total clusters, the hierarchical permutation test allows good control of Type I error rate without regard for the underlying distribution. Even for very asymmetric distributions (such as the power law distribution) type I error rate could be acceptably controlled via hierarchical randomization. For 6 clusters, the hierarchical permutation test performed similarly well, but over-rejects the null for certain distributions when the between-cluster variance is much greater than the within-cluster variance. Using studentized covariance as the test statistic gave similar results (**SI Figure 1**), which is expected as both statistics test for a difference in means.

**Figure 6.**
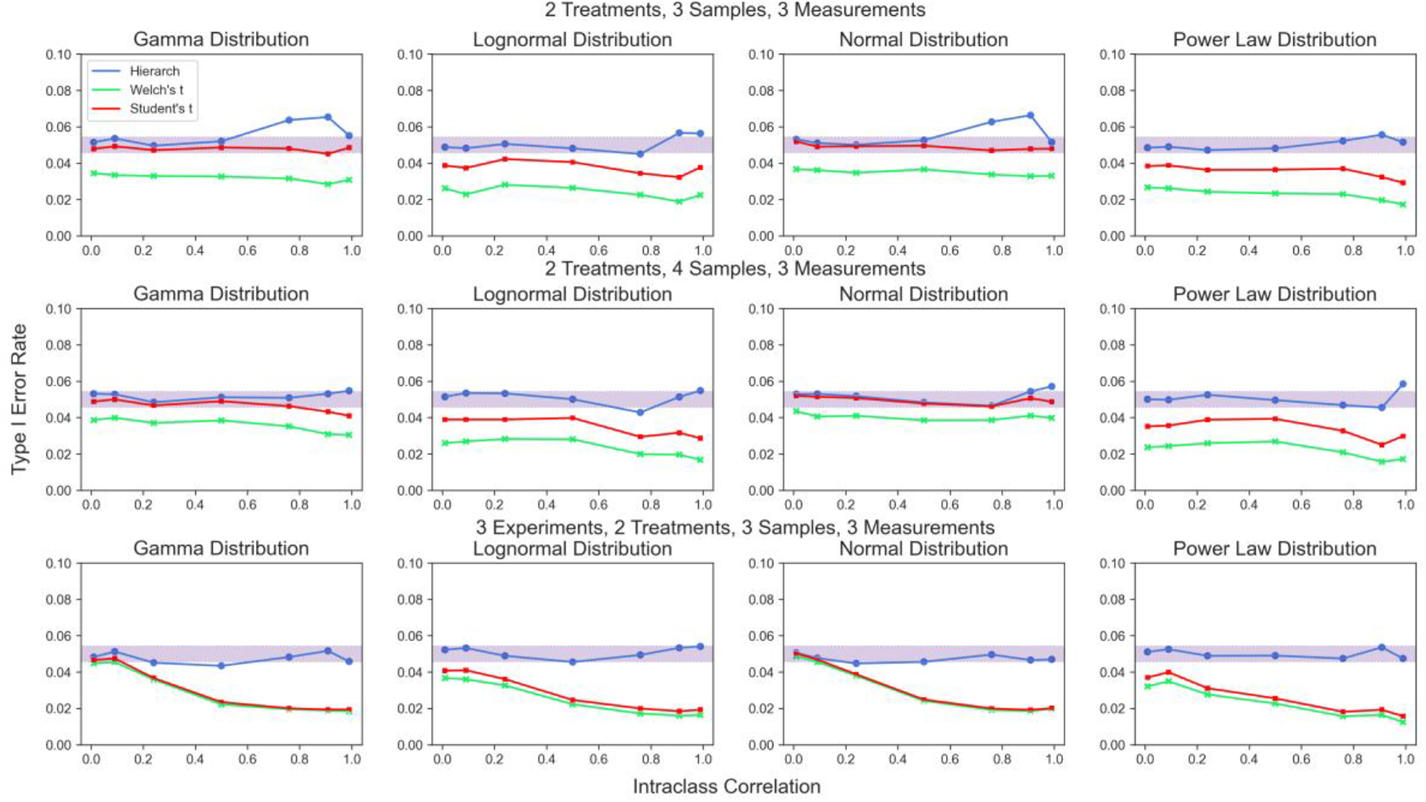
Type I error rate for hierarchical randomization test based on the t statistic compared to Student’s t test and Welch’s t test. Both treatment groups have equal variance. Shaded area represents the 95% binomial confidence interval around a 5% type I error rate.

Next, we investigated Type I error rate in identical conditions as above, but with unequal variances between treatment groups (1:1.5). Again, we found that the hierarchical randomization test afforded better control of Type I error rates than either t-test in most cases, though the performance is slightly worse for very few clusters (**Figure 7**). The usage of a pivotal test statistic gives hierarchical randomization only asymptotic validity against the weak null hypothesis^28^ - as the number of treated samples increases, the better the test performs when the variances of the two samples are different. Again, the performance of the hierarchical randomization test based on studentized covariance was essentially identical to the performance of the hierarchical randomization test based on the t statistic in these conditions (**SI Figure 2**).

**Figure 7.**
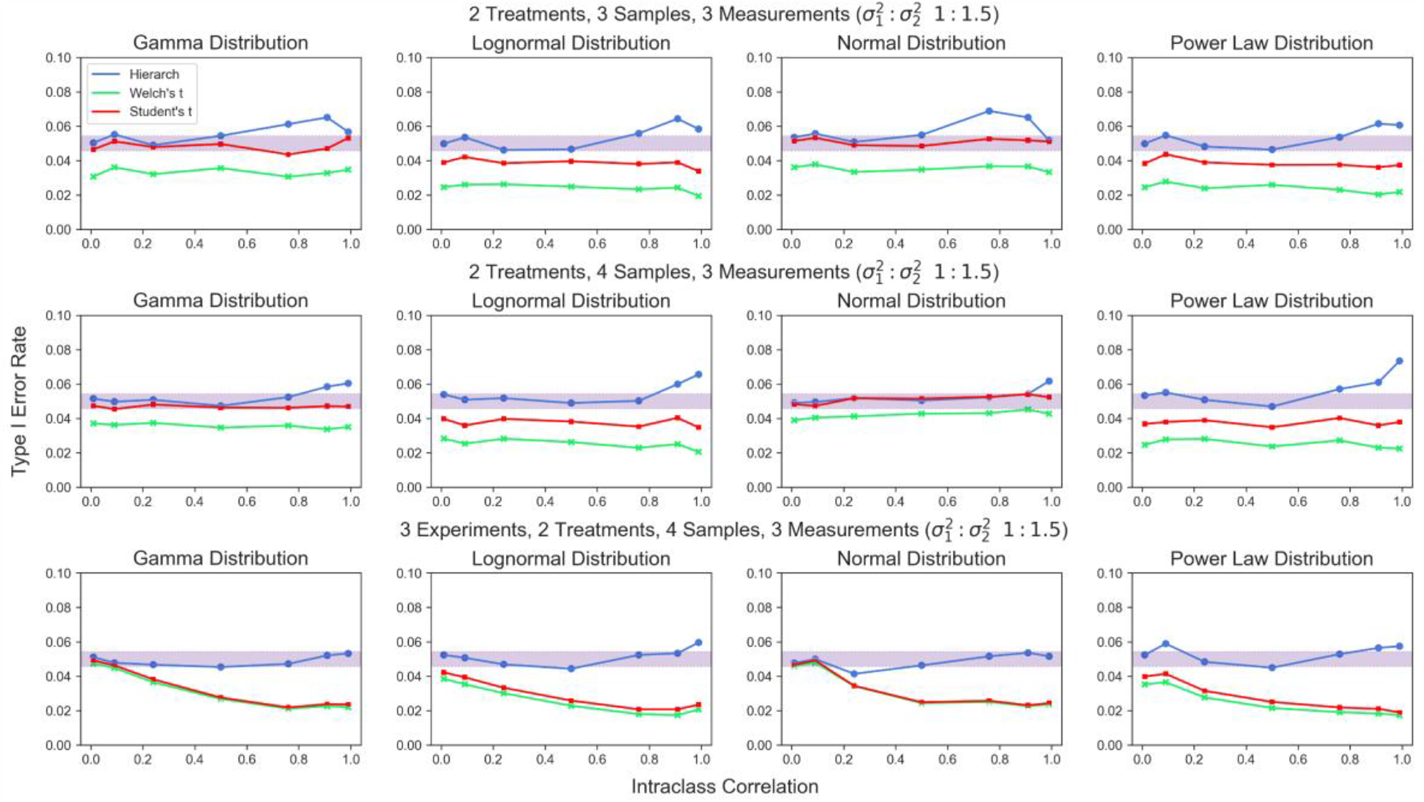
Type I error rate for hierarchical randomization test based on the t statistic, Welch’s t test, and Student’s t test given unequal between-cluster variance. Shaded area represents the 95% binomial confidence interval around a 5% type I error rate.

To validate the performance of the hierarchical randomization test with multiple treatment conditions, we simulated data under identical conditions as above, but with 3 or 4 treatment groups (**Figure 8**). We found that unlike the standard Wald test, hierarchical randomization maintains Type I error rate control when presented with non-normal errors and maintains good control in the presence of heteroscedasticity (**SI Figure 3**).

**Figure 8.**
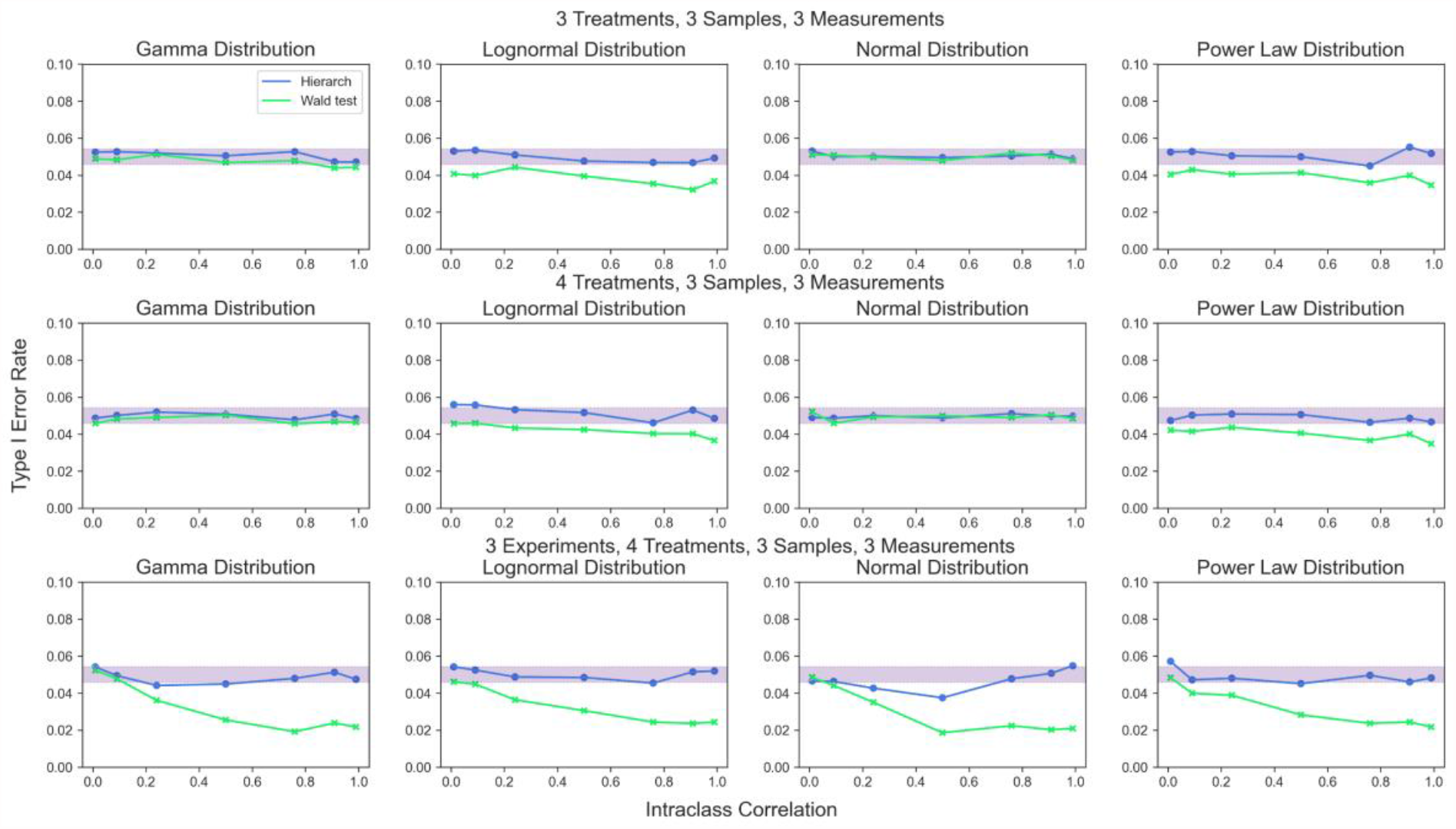
Type I error rate for hierarchical randomization test and the Wald test. Shaded area represents the 95% binomial confidence interval around a 5% type I error rate.

Finally, we examined the confidence intervals generated by hierarchical randomization. We simulated data with a range of mean differences to monitor the coverage probability of 95% confidence intervals computed by hierarchical randomization (**Figure 9**). We found that inverting a hierarchical randomization test produces 95% confidence intervals that maintain nominal coverage without regard for the underlying distribution. The studentized covariance test statistic performs similarly well for datasets with multiple treatment groups (**Figure 10**).

**Figure 9.**
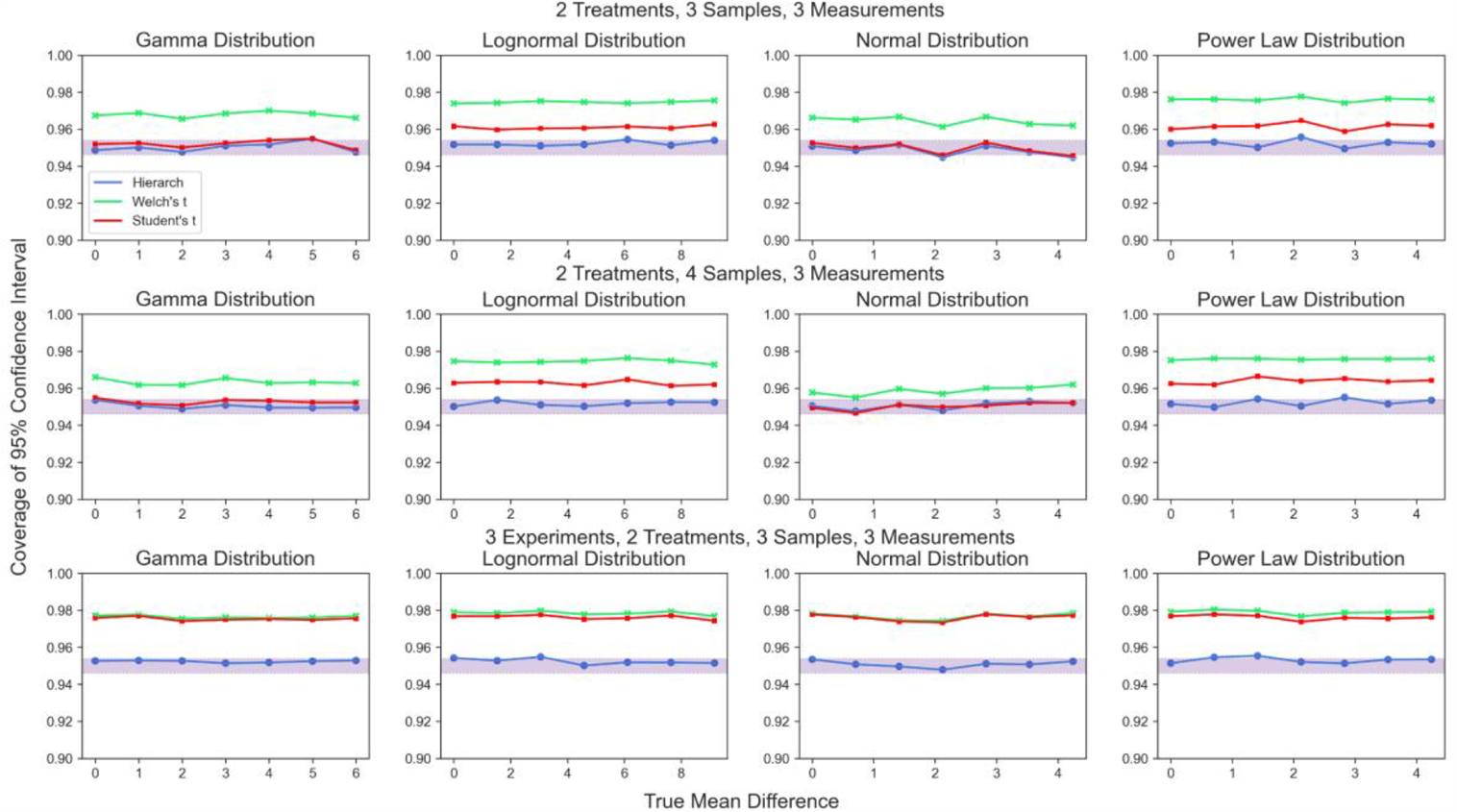
Coverage probability of 95% confidence intervals generated by test inversion. Shaded area represents the 95% binomial confidence interval around a 95% coverage probability.

**Figure 10.**
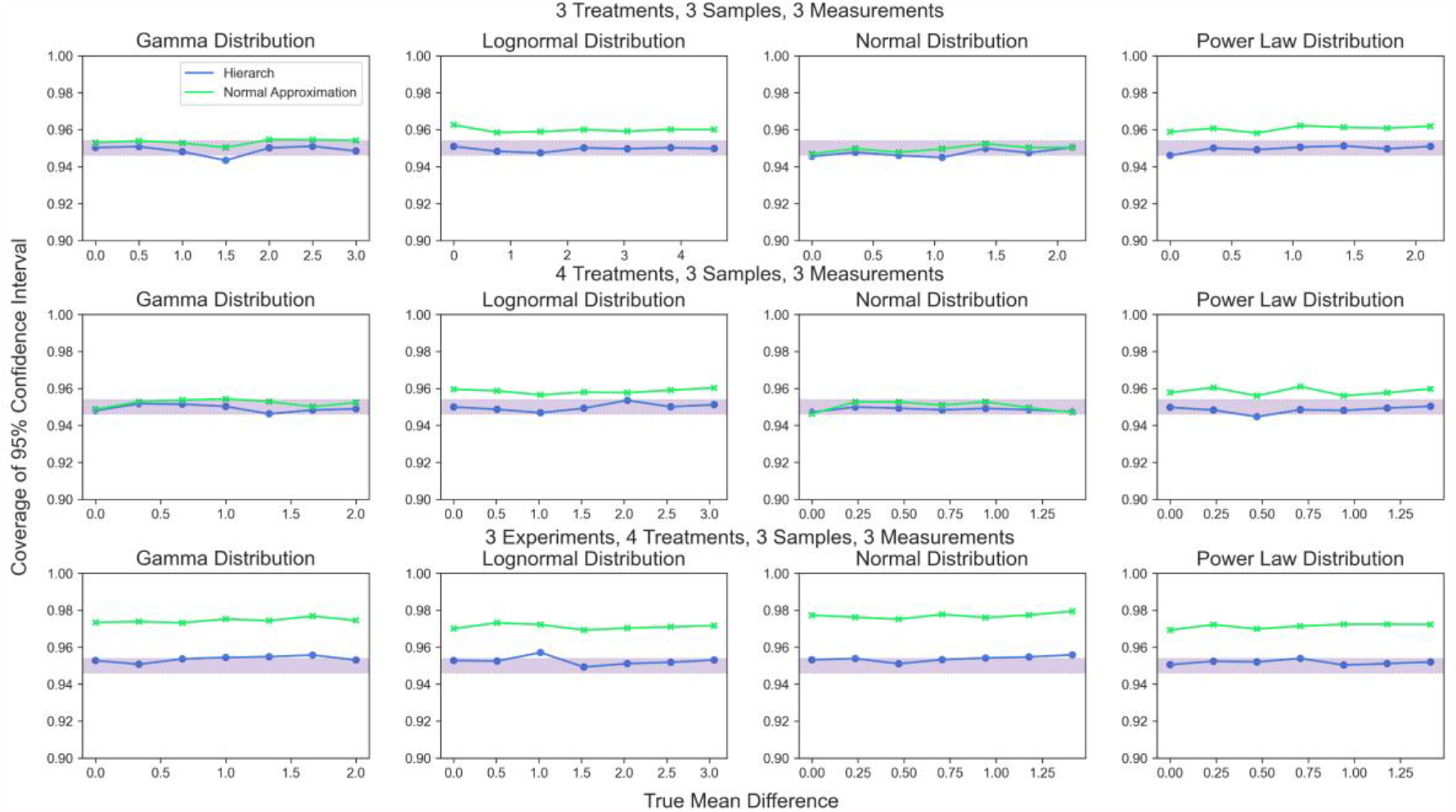
Coverage probability of 95% confidence intervals generated by test inversion. Shaded area represents the 95% binomial confidence interval around a 95% coverage probability.

In short, hierarchical randomization is a flexible strategy for performing hypothesis tests and computing confidence intervals that can handle many levels of data while maintaining robustness to non-normal errors and heteroscedasticity. Furthermore, we find the construction of the test to be pedagogical - despite our reliance on null hypothesis testing, many researchers have only a fuzzy conception of what a p value is telling them. By generating an empirical null distribution by resampling in a manner mimicking the experimental design, hierarchical permutation tests open the black box of hypothesis testing and make the meaning of a two-tailed test and the resulting p value intuitive: taking it as a given that the treatment had no effect, what are all the possible values of the test statistic that can be generated by shuffling the “treatment” labels on the data? What fraction of those possible test statistics are as or more extreme than the observed test statistic?

## Discussion

Hierarchical randomization tests enable researchers to analyze a wide variety of experimental designs while retaining good control of Type I error rate and freedom from distributional assumptions. By using a mix of permuting and bootstrapping, these tests incorporate all of the information in a dataset without unnecessary summarization steps. In this manner, hierarchical randomization tests have an element of a Bayesian approach - rather than summarizing information from lower levels using point estimates, the randomization test is performed on the full posterior distribution of each cluster. We use simulation studies to confirm that this approach controls the two-tailed Type I error rate at 0.05 when there are as few as eight total clusters and when the intra-class correlation is sufficiently low for six clusters, which is impossible for traditional cluster permutation tests.

Despite their excellent statistical properties, randomization tests have historically been restricted to certain subfields.^44–46^ We feel this is in part because setting up a randomization test often requires custom code, which results in a high computational burden and slow analyses. The software package presented in this work, *Hierarch*, makes setup and execution of a hierarchical test much simpler for practicing researchers. Not only can *Hierarch* infer the design of an experiment from the layout of the input data, but it is quite fast - every test described in this work (which use up to 10x the number of permutations necessary to compute a precise p-value) can be performed in under a second on a wide range of personal laptop computers.

We also introduce a covariance-based test statistic based on the work of DiCiccio and Romano to construct a randomization test that can be applied to linear regression problems.^42^ This test statistic is approximately pivotal and, in the two-sample case, is linearly related to the t-statistic. We use simulation studies to confirm that this studentized covariance randomization test retains the t-statistic’s Type I error rate control under homoscedasticity, and also has the desired asymptotic validity under heteroscedasticity. Notably, the test performs well even when the data is drawn from the heavily asymmetric power law distribution under heteroscedastic conditions. Even when the test fails, the Type I error rate is quite close to 5%. We demonstrate that hierarchical randomization tests based on both the t statistic and studentized covariance can be inverted to form effect size confidence intervals that maintain nominal coverage regardless of the underlying distribution.

Hierarchical randomization hypothesis tests are much like the jackknife – they can be applied to a wide variety of experimental designs while maintaining better control of Type I error rates than asymptotic tests. While another statistical test may perform better for datasets that fulfill the assumptions of that test, these assumptions are often unverifiable – hierarchical randomization tests can be applied to any hierarchical dataset and produce an answer that does not depend on unverifiable assumptions. They do this by including multiple levels of clustering without discarding information from any level of the experimental design. These tests have good small-sample properties and are valid for several nested experimental designs common to biological research. In most cases, hypothesis testing with *Hierarch* can achieve the “platinum standard” of significance analysis without requiring researchers to produce custom code or to wait minutes or hours for their computers to produce a *p*-value.

## Code Availability

The *Hierarch* source code is available at https://github.com/rishi-kulkarni/hierarch. The simulations performed in this manuscript are available as Jupyter Notebooks at https://github.com/rishi-kulkarni/hierarch-simulations.

## Acknowledgements

We would like to thank Katherine Derosier, Alex Truong, and Eric Xia for helpful discussions regarding implementation of the algorithm and preparation of the manuscript. This work was generously supported by the US National Institutes of Health (Grant R01 CA227942). Figure illustrations were created using BioRender.com.

## Author Contributions

Conceptualization, RUK and CRB.; Methodology and Formal Analysis, RUK., Investigation, RUK and CLW.; Writing – Original Draft, RUK.; Writing – Review and Editing, CLW and CRB.; Funding Acquisition – CRB.; Supervision, CRB.

## References

1. Galbraith, S., Daniel, J. A. & Vissel, B. A Study of Clustered Data and Approaches to Its Analysis. J. Neurosci. 30, 10601–10608 (2010).

2. Huang, F. L. Using Cluster Bootstrapping to Analyze Nested Data With a Few Clusters. Educ. Psychol. Meas. 78, 297–318 (2018).

3. Moen, E. L., Fricano-Kugler, C. J., Luikart, B. W. & O’Malley, A. J. Analyzing Clustered Data: Why and How to Account for Multiple Observations Nested within a Study Participant? PLOS ONE 11, e0146721 (2016).

4. Saravanan, V., Berman, G. J. & Sober, S. J. Application of the hierarchical bootstrap to multi-level data in neuroscience. ArXiv200707797Q-Bio (2020).

5. Musca, S. C. et al. Data with Hierarchical Structure: Impact of Intraclass Correlation and Sample Size on Type-I Error. Front. Psychol. 2, (2011).

6. Dowding, I. & Haufe, S. Powerful Statistical Inference for Nested Data Using Sufficient Summary Statistics. Front. Hum. Neurosci. 12, 103 (2018).

7. Greenland, S. et al. Statistical tests, P values, confidence intervals, and power: a guide to misinterpretations. Eur. J. Epidemiol. 31, 337–350 (2016).

8. Handbook of multilevel analysis. (Springer, 2008).

9. Leeden, R. van der, Meijer, E. & Busing, F. M. T. A. Resampling Multilevel Models. in Handbook of Multilevel Analysis (eds. Leeuw, J. de & Meijer, E.) 401–433 (Springer, 2008). doi:10.1007/978-0-387-73186-5_11.

10. Maas, C. J. M. & Hox, J. J. Sufficient Sample Sizes for Multilevel Modeling. Methodol. Eur. J. Res. Methods Behav. Soc. Sci. 1, 86–92 (2005).

11. Schielzeth, H. et al. Robustness of linear mixed-effects models to violations of distributional assumptions. Methods Ecol. Evol. 11, 1141–1152 (2020).

12. Austin, P. C. & Leckie, G. The effect of number of clusters and cluster size on statistical power and Type I error rates when testing random effects variance components in multilevel linear and logistic regression models. J. Stat. Comput. Simul. 88, 3151–3163 (2018).

13. Berger, V. W. Pros and cons of permutation tests in clinical trials. Stat. Med. 19, 1319–1328 (2000).

14. Bind, M.-a. C. & Rubin, D. B. When possible, report a Fisher-exact P value and display its underlying null randomization distribution. Proc. Natl. Acad. Sci. 117, 19151–19158 (2020).

15. Bertanha, M. & Chung, E. Permutation Tests at Nonparametric Rates. ArXiv 210213638 Econ Math Stat (2021).

16. Janssen, A. Studentized permutation tests for non-i.i.d. hypotheses and the generalized Behrens-Fisher problem. Stat. Probab. Lett. 36, 9–21 (1997).

17. Tukey, J. W. Tightening the clinical trial. Control. Clin. Trials 14, 266–285 (1993).

18. Berger, V., Lunneborg, C., Ernst, M. & Levine, J. Parametric Analyses In Randomized Clinical Trials. J. Mod. Appl. Stat. Methods 1, (2002).

19. Harris, C. R. et al. Array programming with NumPy. Nature 585, 357–362 (2020).

20. Lam, S. K., Pitrou, A. & Seibert, S. Numba: a LLVM-based Python JIT compiler. in Proceedings of the Second Workshop on the LLVM Compiler Infrastructure in HPC 1–6 (Association for Computing Machinery, 2015). doi:10.1145/2833157.2833162.

21. Abadie, A., Athey, S., Imbens, G. W. & Wooldridge, J. When Should You Adjust Standard Errors for Clustering? http://www.nber.org/papers/w24003 (2017) doi:10.3386/w24003.

22. Winkler, A. M., Webster, M. A., Vidaurre, D., Nichols, T. E. & Smith, S. M. Multi-level block permutation. NeuroImage 123, 253–268 (2015).

23. Efron, B. Bootstrap Methods: Another Look at the Jackknife. Ann. Stat. 7, 1–26 (1979).

24. Becher, H., Hall, P. & Wilson, S. R. Bootstrap Hypothesis Testing Procedures. Biometrics 49, 1268–1272 (1993).

25. Beran, R. Prepivoting Test Statistics: A Bootstrap View of Asymptotic Refinements. J. Am. Stat. Assoc. 83, 687–697 (1988).

26. Rubin, D. B. The Bayesian Bootstrap. Ann. Stat. 9, 130–134 (1981).

27. Breiman, L. Bagging predictors. Mach. Learn. 24, 123–140 (1996).

28. Chung, E. & Romano, J. P. Exact and asymptotically robust permutation tests. Ann. Stat. 41, 484–507 (2013).

29. Wasserstein, R. L. & Lazar, N. A. The ASA Statement on p-Values: Context, Process, and Purpose. Am. Stat. 70, 129–133 (2016).

30. Dunkler, D., Haller, M., Oberbauer, R. & Heinze, G. To test or to estimate? P-values versus effect sizes. Transpl. Int. 33, 50–55 (2020).

31. Falk, M. & Kaufmann, E. Coverage Probabilities of Bootstrap-Confidence Intervals for Quantiles. Ann. Stat. 19, 485–495 (1991).

32. Hall, P. On the Number of Bootstrap Simulations Required to Construct a Confidence Interval. Ann. Stat. 14, 1453–1462 (1986).

33. Algina, J., Keselman, H. J. & Penfield, R. D. Confidence Interval Coverage for Cohen’s Effect Size Statistic. Educ. Psychol. Meas. 66, 945–960 (2006).

34. Loughin, T. M. A systematic comparison of methods for combining p-values from independent tests. Comput. Stat. Data Anal. 47, 467–485 (2004).

35. Zucker, D. M., Lieberman, O. & Manor, O. Improved Small Sample Inference in the Mixed Linear Model: Bartlett Correction and Adjusted Likelihood. J. R. Stat. Soc. Ser. B Stat. Methodol. 62, 827–838 (2000).

36. Luke, S. G. Evaluating significance in linear mixed-effects models in R. Behav. Res. Methods 49, 1494–1502 (2017).

37. Winkler, A. M., Ridgway, G. R., Webster, M. A., Smith, S. M. & Nichols, T. E. Permutation inference for the general linear model. NeuroImage 92, 381–397 (2014).

38. Riley, R. D., Higgins, J. P. T. & Deeks, J. J. Interpretation of random effects meta-analyses. BMJ 342, d549 (2011).

39. Konietschke, F. & Pauly, M. A studentized permutation test for the nonparametric Behrens-Fisher problem in paired data. Electron. J. Stat. 6, 1358–1372 (2012).

40. Neubert, K. & Brunner, E. A studentized permutation test for the non-parametric Behrens– Fisher problem. Comput. Stat. Data Anal. 51, 5192–5204 (2007).

41. Janssen, A. & Pauls, T. A Monte Carlo comparison of studentized bootstrap and permutation tests for heteroscedastic two-sample problems. Comput. Stat. 20, 369–383 (2005).

42. DiCiccio, C. J. & Romano, J. P. Robust Permutation Tests For Correlation And Regression Coefficients. J. Am. Stat. Assoc. 112, 1211–1220 (2017).

43. Manly, B. F. J. Randomization, Bootstrap and Monte Carlo Methods in Biology. (Chapman and Hall/CRC, 2017). doi:10.1201/9781315273075.

44. Ludbrook, J. & Dudley, H. Why Permutation Tests Are Superior to t and F Tests in Biomedical Research. Am. Stat. 52, 127–132 (1998).

45. Doerge, R. W. & Churchill, G. A. Permutation Tests for Multiple Loci Affecting a Quantitative Character. Genetics 142, 285–294 (1996).

46. Wen, B. et al. IQuant: An automated pipeline for quantitative proteomics based upon isobaric tags. PROTEOMICS 14, 2280–2285 (2014).

