## Supplemental Information for "Analyzing Nested Experimental Designs – A User-Friendly Resampling Method to Determine Experimental Significance"

**SI Appendix 1**

The unbiased estimator for Var(Q) is as follows, where s_r,t_ is the cross power sum of X and Y:

$$S^{2}\left( Q \right)= \frac{1}{\left( -3+n \right)\left( -2+n \right)\left( -1+n \right)^{2}n^{2}} \left( \left( 6-4n \right)s_{0,1}^{2}s_{1,0}^{2}+\left( -n+n^{2} \right)s_{0,2}s_{1,0}^{2}+\left( -10n+6n^{2} \right)s_{0,1}s_{1,0}s_{1,1}+\left( 2n+n^{2}-n^{3} \right)s_{1,1}^{2}+\left( -2n+4n^{2}-2n^{3} \right)s_{1,0}s_{1,2}+\left( -n+n^{2} \right)s_{0,1}^{2}s_{2,0}+\left( n-n^{2} \right)s_{0,2}s_{2,0}+\left( -2n+4n^{2}-2n^{3} \right)s_{0,1}s_{2,1}+\left( n^{2}-2n^{3}+n^{4} \right)s_{2,2} \right)$$

$$s_{r,t}= \sum_{i=0}^{n} \left( X_{i} \right)^{r}\left( Y_{i} \right)^{t}$$

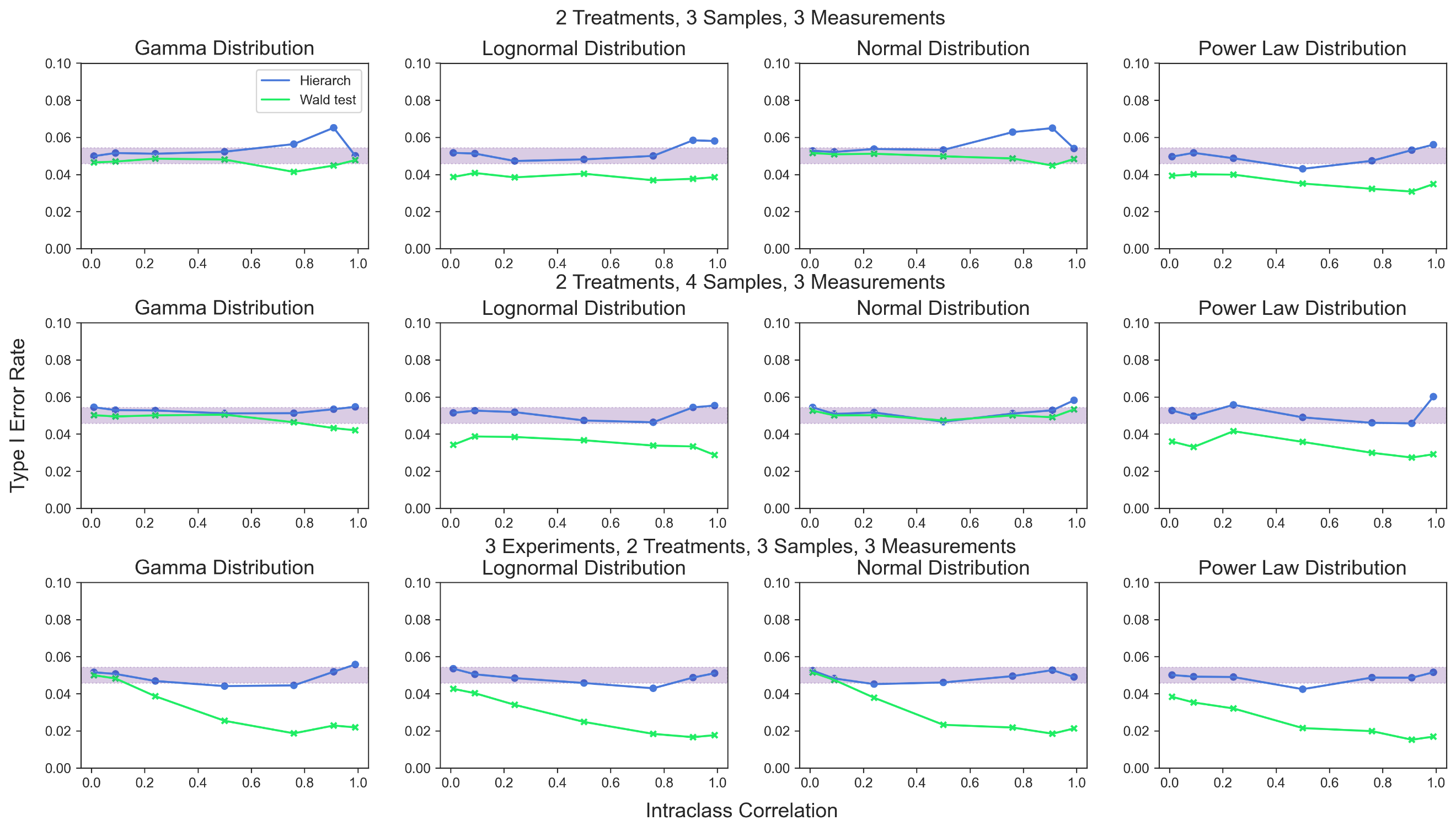

**SI Figure 1. Type I error rate for hierarchical randomization test based on studentized covariance and the Wald test.** Shaded area represents the 95% binomial confidence interval around a 5% type I error rate.

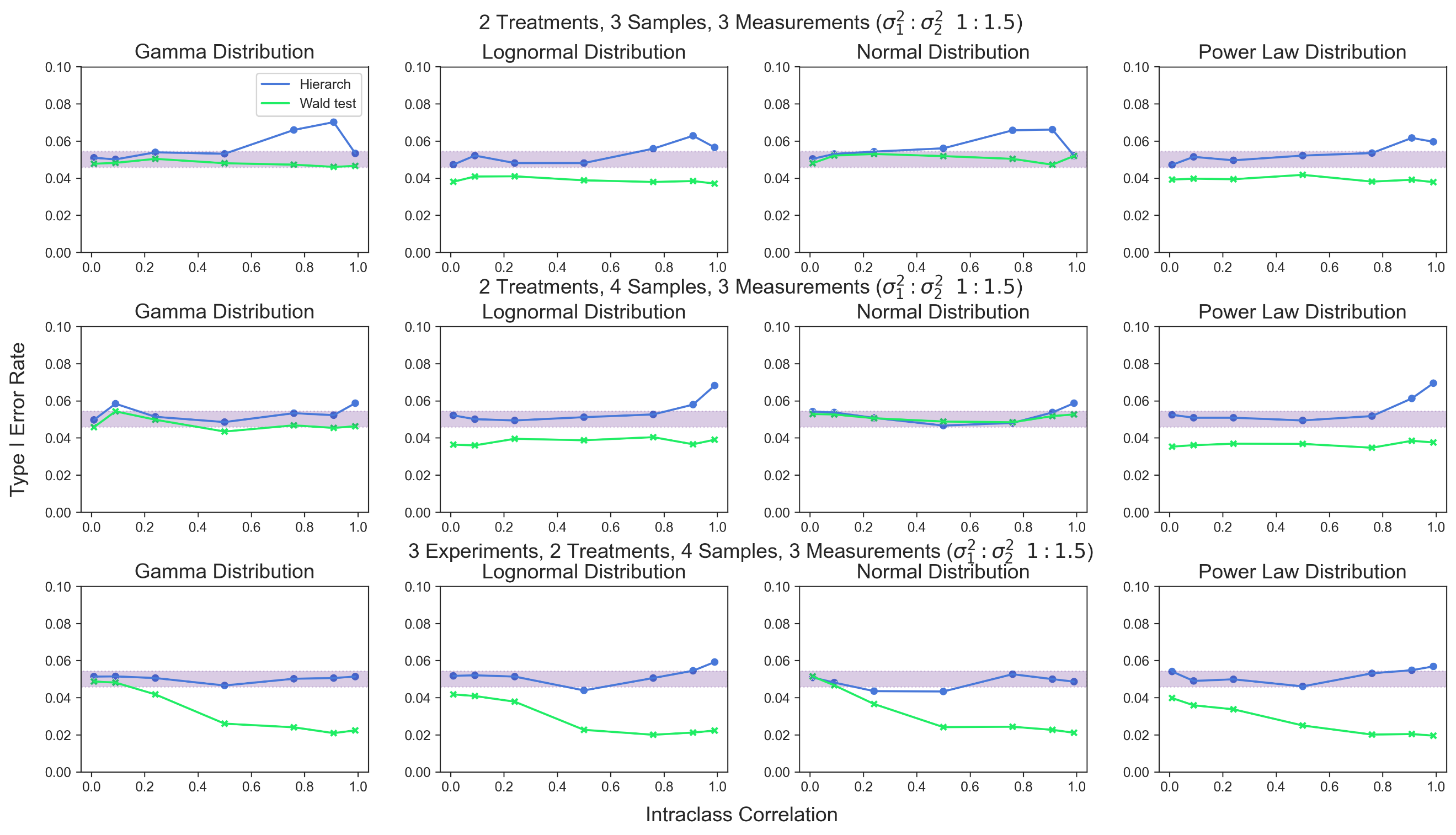

**SI Figure 2. Type I error rate for hierarchical randomization test based on studentized covariance and the Wald test given unequal between-cluster variance.** Shaded area represents the 95% binomial confidence interval around a 5% type I error rate.

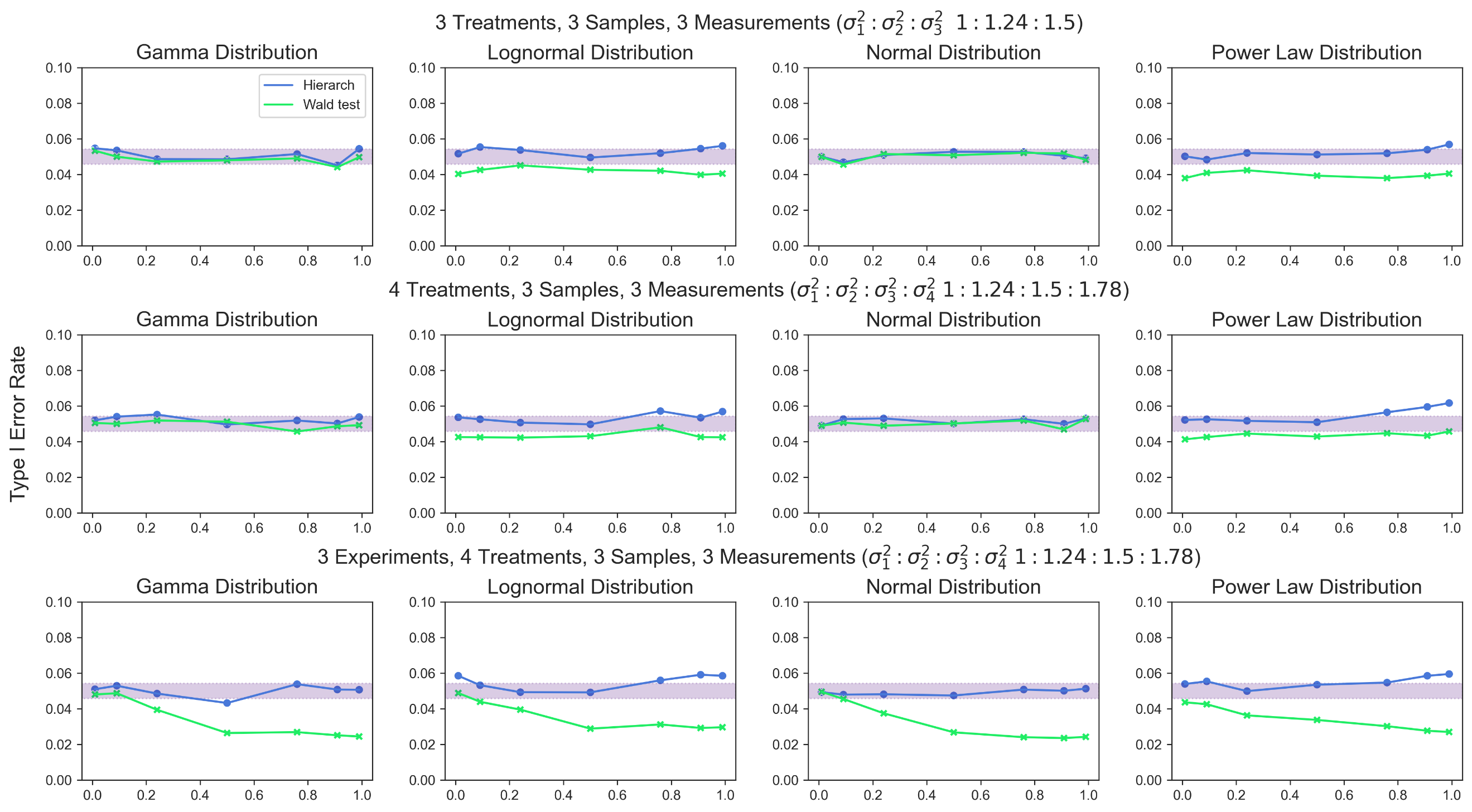

**SI Figure 3. Type I error rate for hierarchical randomization test and the Wald test given unequal variances.** Shaded area represents the 95% binomial confidence interval around a 5% type I error rate.
